# Improved long-read sequencing technology reveals full genome architecture of Malian *Plasmodium falciparum* field isolates

**DOI:** 10.64898/2026.11.13.766388

**Authors:** B. Díaz-Terenti, M. Diakite, L. C. Ranford-Cartwright, S. Pradhan, M.C. Martos-Contreras, T. D. Otto, E. Gómez-Díaz

**Affiliations:** Instituto de Parasitología y Biomedicina López-Neyra (IPBLN), Consejo Superior de Investigaciones Científicas, 18016, Granada, Spain; School of Biodiversity, One Health and Veterinary Medicine, College of Medical, Veterinary and Life Sciences, University of Glasgow, Glasgow, G12 8QQ, UK; University of Science, Techniques, and Technologies of Bamako, Mali; Centre of Immunobiology, Institute of Infection, Immunity & Inflammation, College of Medical, Veterinary and Life Sciences University of Glasgow, Glasgow, UK; Bernhard Nocht Institute for Tropical Medicine (BNITM); Universität Hamburg (UHH)

**Keywords:** Malaria parasites, PacBio HiFi, de novo genome assembly, natural genomic diversity, subtelomeric regions, telomeres, multigene families

## Abstract

Understanding genome plasticity in malaria parasites and the diversity of their repetitive gene families requires high-quality reference genomes derived from parasites recently isolated from patients. However, generating such genomes in *Plasmodium falciparum* remains challenging owing to the parasite’s extreme AT content, extensive repetitive regions, and the frequent occurrence of multiclonal infections in endemic settings. As a result, most available reference genomes are derived from laboratory-adapted strains, that have experienced prolonged *in vitro* maintenance, and that incompletely represent natural parasite diversity.

We developed and optimized a *de novo* assembly strategy based on PacBio HiFi long-read sequencing and applied it to eight recently-isolated and cloned *P. falciparum* parasites from Mali. HiFi sequencing enabled highly accurate genome reconstruction, yielding assemblies with a mean of 18 contigs, 301 pseudogenes, and a frameshift rate of 0.04% following Liftoff annotation. Assemblies showed high structural integrity and near-complete chromosomes, including improved recovery of subtelomeric regions and multigene families, without requiring short-read correction. Despite overall conservation of chromosome organization, substantial variation was observed in chromosome-end architecture, subtelomeric structure, and telomeric repeat content among parasite genomes.

Our approach simplifies the generation of high-quality *P. falciparum* genomes and reduces sequencing requirements fourfold. The resulting assemblies enabled characterization of natural variation in telomere plasticity and multicopy gene family diversity in field isolates. These findings establish PacBio HiFi sequencing as an effective and scalable strategy for producing highly contiguous and accurate *de novo* assemblies of *P. falciparum*, supporting the development of geographically-diverse, field-relevant malaria parasite reference genomes.

## Introduction

The human malaria parasite *Plasmodium falciparum* exhibits remarkable phenotypic diversity, which underpins its adaptability and pathogenicity. This diversity, driven by extensive genomic variation arising from mechanisms such as genetic recombination, gene conversion, and duplication, poses significant challenges to malaria control and eradication efforts. The parasite’s unique population dynamics further contribute to this genomic complexity, complicating the development of effective interventions ^1–3^.

Despite the availability of genomic data from more than 20,000 *P. falciparum* field isolates, most datasets rely on short-read sequencing, limiting the resolution of subtelomeric regions and large structural variants^4,5^. In contrast, the few high-quality long-read reference genomes available are predominantly derived from laboratory-adapted or long-term cultured parasites that may not accurately capture the genomic plasticity of natural populations^6^. Bridging this gap requires the generation of representative chromosome-level genome assemblies from recently adapted parasite isolates that capture the genetic diversity circulating in malaria-endemic regions.

However, the *P. falciparum* genome presents several intrinsic challenges that hinder comprehensive sequencing and annotation. Notably, it exhibits an extreme AT bias—the highest known among all organisms—which results in uneven sequencing coverage, increased error rates, and assembly gaps^7^. Additionally, the genome contains extensive low-complexity repetitive regions, particularly in subtelomeric areas, which complicate assembly and obscure functional element identification^5^. The presence of large, highly polymorphic multigene families such as *var*, *rifin*, and *stevor*, often organized in subtelomeric regions, further limits accurate genome representation. These gene families encode proteins critical for immune evasion and host interaction, with *var* genes encoding the PfEMP1 protein that mediates cytoadherence and contributes to severe malaria manifestations^8^. Their complex regulation, including mutually exclusive monoallelic expression, adds another layer of genomic intricacy^9^.

Traditional long-read sequencing technologies, such as Pacific Biosciences (PacBio) and Oxford Nanopore Technologies (ONT), have been instrumental in overcoming some of these challenges by enabling de novo assembly of the *P. falciparum* genome. However, the early application of long-read sequencing to *P. falciparum* was associated with very high error rates, necessitating consensus polishing with Illumina short-read data to distinguish true variants from sequencing errors^10^, and careful assembly strategies to resolve highly repetitive regions. Importantly, successful resolution of subtelomeric regions and recent duplications also depends on obtaining reads long enough to span homologous sequences, with reads >15 kb shown to facilitate telomere-to-telomere genome assembly^5,11^.

The recent advent of PacBio’s high-fidelity (HiFi) reads offers a transformative opportunity for sequencing complex genomes like that of *P. falciparum*. HiFi reads, generated by consensus sequencing of multiple passes over the same DNA molecule, combine the length advantages of long reads with accuracy levels typically associated with short reads (error rates below 1%). This technology simplifies workflows, enhances assembly quality, and reduces costs by minimizing reliance on complementary sequencing approaches and lowering data requirements.

Recent studies have demonstrated the potential of PacBio HiFi sequencing to generate highly contiguous chromosome-scale *P. falciparum* assemblies, improving resolution of subtelomeric regions. HiFi technology has also been applied to targeted sequencing of highly polymorphic antigen genes, enabling full-length haplotype reconstruction and more accurate estimation of infection complexity^12–14^.

Despite these advances, HiFi-based assemblies remain available for only a limited number of field isolates, and globally representative reference genomes are still lacking. Near telomere-to-telomere assemblies remain uncommon, particularly for highly repetitive chromosome ends and structurally complex subtelomeric regions. As a result, key features of parasite genome biology remain poorly understood, including the architecture of variant gene families and the distribution of repetitive elements such as rep20, which are thought to contribute to genome stability, chromosome-end dynamics, recombination, and the regulation of virulence-associated genes^15–17^. Moreover, standardized and cost-effective workflows for routinely generating high-quality de novo assemblies at scale have yet to be established, and the sequencing requirements needed to produce near-complete genomes remain poorly defined.

High-quality chromosome-scale assemblies also have important applications for malaria genomic surveillance. Whole-genome sequencing is increasingly used to monitor the emergence and spread of antimalarial drug resistance, including variants in *pfk13*, *pfcrt*, *pfmdr1*, *pfdhfr*, and *pfdhps*, that directly influence treatment efficacy and malaria control efforts^18,19^. Accurate assembly of field-derived genomes therefore provides an opportunity to simultaneously characterize resistance-associated loci and broader patterns of genome structural variation within contemporary parasite populations.

In this study, we leverage the advanced capabilities of PacBio HiFi sequencing to optimize a novel, comprehensive strategy for the de novo assembly of the *P. falciparum* genome. Our approach aims to simplify the assembly process while improving accuracy and completeness, particularly in challenging genomic regions such as subtelomeric multigene families. This work seeks to generate higher-quality genome assemblies that will facilitate a deeper understanding of antigenic variation, virulence, and parasite-host interactions, ultimately supporting the development of more effective malaria control strategies.

## MATERIALS AND METHODS

### 1. Ethical statement

Clinical isolates used in this study were obtained from malaria-infected individuals enrolled in studies conducted in Mali. All samples were collected under approved ethical protocols, and informed consent or assent was obtained from all participants or their legal guardians prior to sample collection. Ethical approval: Comité d’Ethique Faculté des Facultés de Médecine/d’Odontostomatologie et the Pharmacie (FMOS/FAPH), approval number [2020/227/CE/FMOS/FAPH]; Comité Coordinador de Ética de la Investigación Biomédica de Andalucía, approval number [PEIBA: 2419-N-20]; MVLS Ethics Committee University of Glasgow, approval number [200200074].

### 2. Parasite culture and DNA extraction

The samples used in this study correspond to eight *P. falciparum* clones that were obtained from eight Malian clinical isolates, some of which were polyclonal infections. Full details of the parasite isolates and cloning are given in Pradhan et al (manuscript in preparation*). For genomic DNA extraction and sequencing, parasites were thawed from cryopreserved stocks and cultured in 5 mL flasks for the minimum number of generations necessary to achieve a late-stage parasitaemia between 2% and 4%. Culture was performed in AB+ erythrocytes (5% hematocrit) under standard conditions^20^ with media containing 0.5% Albumax II at 37 □, under a gas mixture of 96% nitrogen, 3% carbon dioxide and 1% oxygen. Genomic DNA was extracted from synchronized trophozoite-stage *P. falciparum* cultures using a modified phenol-chloroform protocol^21^, including SDS lysis, Proteinase K digestion, organic extraction, and isopropanol precipitation of high-molecular-weight DNA. DNA quality and concentration were determined by spectrophotometry (A260/280) and 1% agarose gel electrophoresis (Supplementary Table S1).

### 3. PacBio HiFi sequencing

The eight clones were sequenced using *Pacific Bioscience (PacBio) real-time sequencing technology* (single molecule real time, SMRT) to obtain HiFi Long reads with high accuracy (99.9%) and an average length of 17 kb. Library preparation and sequencing was outsourced (Centre for Genomic Research, University of Liverpool). Genomic DNA, purity and concentration were quantified using a Qubit fluorometer (Thermo Fisher Scientific). At least 5ug of gDNA per library, obtained from 4-5 flask at 2-4% of parasitemia, was used. DNA libraries were generated using the standard protocol for the Sequel II Binding Kit 3.2 (PacBio). This included DNA fragmentation into fragments of ∼15-20 kb, end repair, ligation of adaptors and library purification. The prepared libraries were subjected to polymerase binding using the Sequel II Binding Kit 3.2. The sequencing of HiFi reads was performed using PacBio technology on a Sequel II system, following the manufacturer’s recommendations. After sequencing, the raw data were subjected to quality control using PacBio tools, checking key metrics such as average read length, consensus accuracy and expected coverage. All resulting information is in Supplementary Table S1. For comparison between conventional PacBio long reads and the PacBio HiFi long reads, we used external data by *Vembar et al. 2016* ^11^ corresponding to conventional PacBio long reads from the 3D7 reference clone of *P. falciparum*.

### 4. Illumina sequencing

Whole genome Illumina sequencing was performed for two of the clones (KK2403 and BG1701) using a PCR-free DNA library preparation kit, designed to minimise PCR-induced bias and provide uniform coverage across the genome. The prepared libraries were then quantified and pooled for sequencing on the Illumina NovaSeq 6000 platform, configured for 2 × 150 bp paired-end reads^22^. Sequencing information is in Supplementary Table S1.

### 5. PacBio HiFi data analyses

#### 4.1. *De novo* genome assembly of new *P. falciparum* clones

*De novo* genome assembly of the 8 newly generated *P. falciparum* clones, was carried out using *HiCanu*, a modified version of the Canu assembler optimized for PacBio HiFi reads, to construct initial draft assemblies. *HiCanu* utilizes homopolymer compression and overlap-based error correction to achieve high accuracy in assembling long reads, which is particularly effective for resolving segmental duplications and complex genomic regions^23^.

All parameters were set at their default values with the following exceptions: (i) the genome size was set at 24,000,000 bp. (ii) the maximum error rate of overlap for correction (*corMaxEvidenceErate*) was set at 0.15 which is the value recommended for high-AT/GC genomes as is the case for *P. falciparum*. *HiCanu* v2.2 was run with the coverage parameter set to 200X, resulting in random selection of reads from the input dataset to achieve the specified coverage.

Following the initial assembly, the *Improvement of Long Read Assemblies (ILRA)* pipeline was employed to enhance the genome assembly by refining contig continuity and accuracy^10^. In this case the parameters of *ILRA* used were *-C no -L pb -K yes -M 245g* avoiding the correction step. Subsequently, *Pilon*^24^ was used to polish the assembly executing one iteration with the assembly resulting from *ILRA*. Unless otherwise stated we used *P. falciparum* 3D7 reference genome for comparisons (release 63 in PlasmoDB)^7^.

*Companion*^25^ was used to annotate the genome assembly. The AUGUSTUS score threshold was set at 0.3. The minimum match length for contig placement ranged from 5000 to 8050, depending on the sample. Gene annotation transfer was performed using two approaches: (i) *Liftoff*^26^ to transfer reference gene models, and (ii) *Rapid Annotation Transfer Tool (RATT)*^27^ with the Assembly transfer type to transfer reference gene models. The annotations generated by both methods were compared in terms of annotation quality (see below). Both methods are expected to give similar results for genomes that closely match their reference^25^.

We used the percentage of frameshifts between the *de novo* assemblies and the reference genome at orthologous genes as a measure of annotation quality (given by *Companion*). For this we built a custom script comparing the number of exons in our assembly and the number of exons in the *gff* of the reference genome (script called *Frameshifts_detection.R*). A low number of frameshifts indicates high annotation accuracy, whereas a high number may reflect assembly or annotation issues.

Finally, in *P. falciparum* it is necessary to check the coverage for the hypervariable regions in each sample. In this study, we examined the number of annotated *var, rifin and stevor* genes. We identified LARS motifs in *var* genes using a custom script (getMotifCount.sh). LARS motifs, and motifs associated with *rifin* and *stevor* genes were then used in several subsequent analyses.

The genomes were aligned against the 3D7 reference genome using *minimap2* to assess synteny and structural variations using SyRI, and genomic collinearity was visualized using the *R* package *circlize*^28–30^. The distribution and count of telomeric repeats at the chromosome ends were quantified using *tidk*^31^ with a window size of 5000.

#### 4.2. Comparison of *PacBio* HiFi and original reads with and without Illumina read correction

We define two different steps in the workflow, the first is after the assembly with *HiCanu*, executed as described above. The second step corresponds to the final assembly, after the polishing with *Pilon* in the case of HiFi reads, and after the correction with short reads which is done with *ILRA* in the case of PacBio original.

*HiCanu* assembly was applied to external *P. falciparum* 3D7 conventional PacBio data from Vembar *et al.* (2016)^11^. This was followed by correction of Illumina reads with *ILRA* in combination with *Pilon*. The parameters used were *-C yes -I illumina.fastq -L pb -K yes -Mj 245g -p yes* testing both the default three iterations and five iterations using the -i option.

For two samples (KK2403 and BG1701) for which we had PacBio HiFi and Illumina data, the effect of skipping the correction step in HiFi reads on assembly quality was evaluated using *Pilon* within *ILRA* replacing the previous *Pilon*-only polishing step.

#### 4.3. Coverage reduction with PacBio HiFi reads

The depth required for a *de novo* genome sequencing study depends on the error rate of the sequencing method, the assembly algorithms used, the repetitive complexity of the genome under study and the read length^32^.

To investigate the effect of sequencing depth on genome assembly quality and establish a minimum PacBio HiFi coverage threshold, we used the Lander-Waterman equation (1988) to calculate nominal genome coverage, defined as C=LN/GC, where L is the read length, N is the number of reads, and G is the haploid genome size. Although this metric is widely used to quantify sequencing depth, it assumes a random distribution of reads across the genome and does not account for factors that influence long-read assembly performance, such as uneven coverage, repetitive regions, or structural variation. We therefore used the Lander-Waterman equation solely to standardize coverage estimates, while assessing the impact of coverage on assembly quality empirically through downstream assembly metrics^32^.

To evaluate the effect of sequencing depth on assembly quality, PacBio HiFi datasets were subsampled to target coverages of 5X, 10X, 25X, 50X, 100X, and 200X. The upper limit of 200X was selected because it corresponds to the default coverage parameter used by the *HiCanu* assembler (v2.2). For the dataset from Vembar et al. (2016)^11^, an additional comparison was performed between the original sequencing depth (450X) and a subsampled dataset at 200X coverage.

To obtain the synthetic raw reads we performed a random extraction with *seqtk* (https://github.com/lh3/seqtk) specifying the percentage of the total reads necessary to obtain the desired depth. This method was applied to the HiFi reads for all 6 field clones, and to the conventional PacBio reads for the 3D7 sample.

For each coverage test we applied the same genome assembly pipeline described in the previous section. The assemblies obtained were evaluated in terms of quality and completeness, allowing the identification of the minimum threshold necessary for accurate genome reconstruction.

### 5. Resistant drugs mutation in PacBio Assemblies

Proteome and genome of *Plasmodium falciparum* 3D7 release 63 were retrieved downloaded from PlasmoDB. A *tblastn* search (v2.5.0) of the resistant mutated proteins was performed against each assembly using an E-value threshold of 1e-20, percentage identity >= 80% and coverage >= 85%. The best hit of each resistance-associated protein was selected based on the highest bit score. Genomic regions spanning the hit and 20kb upstream and downstream sequences were extracted for inspection. Protein-to -genome alignment was generated with *Miniprot* (v0.18-r281)^33^ and coding sequences were reconstructed from the resulting gff annotation with *gffread* (v0.12.9)^34^ and *seqkit* (v2.10.0)^35^. Predicted proteins were compared with the corresponding 3D7 protein reference to identify the protein substitutions with a generated R script (*resistant_mutation_identification.R*) using a database of resistance biomarkers. Microsatellite polymorphisms were evaluated separately.

### 6. Statistical analyses

Non-parametric statistical tests were used throughout because of the small sample sizes and repeated-measures study design. Paired comparisons between annotation methods, assembly correction strategies, and polishing iterations were performed using Wilcoxon signed-rank tests, with effect sizes reported as rank-biserial correlations. The effects of sequencing coverage on assembly and annotation metrics were assessed using Friedman tests, with Kendall’s coefficient of concordance (W) used to estimate effect size. Comparisons of multigene-family counts between Malian isolates and reference genomes were conducted using Wilcoxon rank-sum tests. Where multiple related tests were performed, P-values were adjusted using the Benjamini-Hochberg method. All tests were two-sided and statistical significance was assessed at α = 0.05.

## RESULTS

### 1. Near telomere-to-telomere *de novo* genome assemblies of *P. falciparum* using PacBio HiFi long reads

Our assembly pipeline, optimized for PacBio HiFi long reads, generated high-quality assemblies for all eight field-derived clones from Mali, as assessed by multiple assembly metrics (Table 1, Figure S1A). The assemblies had a mean size around 23.5 Mb, closely matching the 23.6-Mb 3D7 reference genome, indicating accurate reconstruction of overall genome content. Assemblies also showed high contiguity, with a mean N50 of 1,672,831 bp (Table 1). The N50 represents the minimum contig length required for contigs of that size or larger to collectively account for at least 50% of the assembled genome; the high N50 values observed therefore indicate substantial assembly continuity. Following *ILRA* and *Pilon* polishing, the number of contigs ranged from 16 to 20 per assembly (Table 1). Given that the *P. falciparum* genome comprises 16 chromosomes (14 nuclear, one mitochondrial and one apicoplast), these values are consistent with highly complete assemblies, while assemblies containing more than 16 contigs likely reflect residual fragmentation. A single 100 bp gap was identified in chromosome 8 of clone BG0801, while two gaps were identified in clone KK2502 on chromosomes 8 and 12 (Table 1). To further assess assembly completeness, we examined chromosome-end structure using the classification scheme shown in Figures 1A-B which is based on the presence or absence of telomeric and subtelomeric features (Fig. 1C, Supplementary Table S2). Across all chromosome ends analysed of the eight de novo assemblies, 74% were classified as complete (State A) In contrast, 13% were classified as State B, lacking the rep20 region but retaining telomeric repeats, while 8.7% were classified as State C, lacking telomeric repeats despite an otherwise complete subtelomeric structure. Only three chromosome ends, one each in clones KK2504, BG0801, and BG1905, were classified as incomplete (State D), representing 3.6% of all chromosome ends analysed. One assembly, KK2403, exhibited a distinct chromosome-end profile compared with the other isolates. Of its 28 chromosome ends, 15 (53.5%) were complete (State A) and 11 (39.3%) belonged to State B. Only two ends lacked telomeric repeats (State C), and no ends were classified as incomplete (State D). Although the high proportion of State B chromosome ends initially suggested reduced completeness, inspection of the underlying HiFi reads strongly supported these structures, indicating that they are unlikely to result from assembly artifacts or sequencing errors.

**Figure 1.**
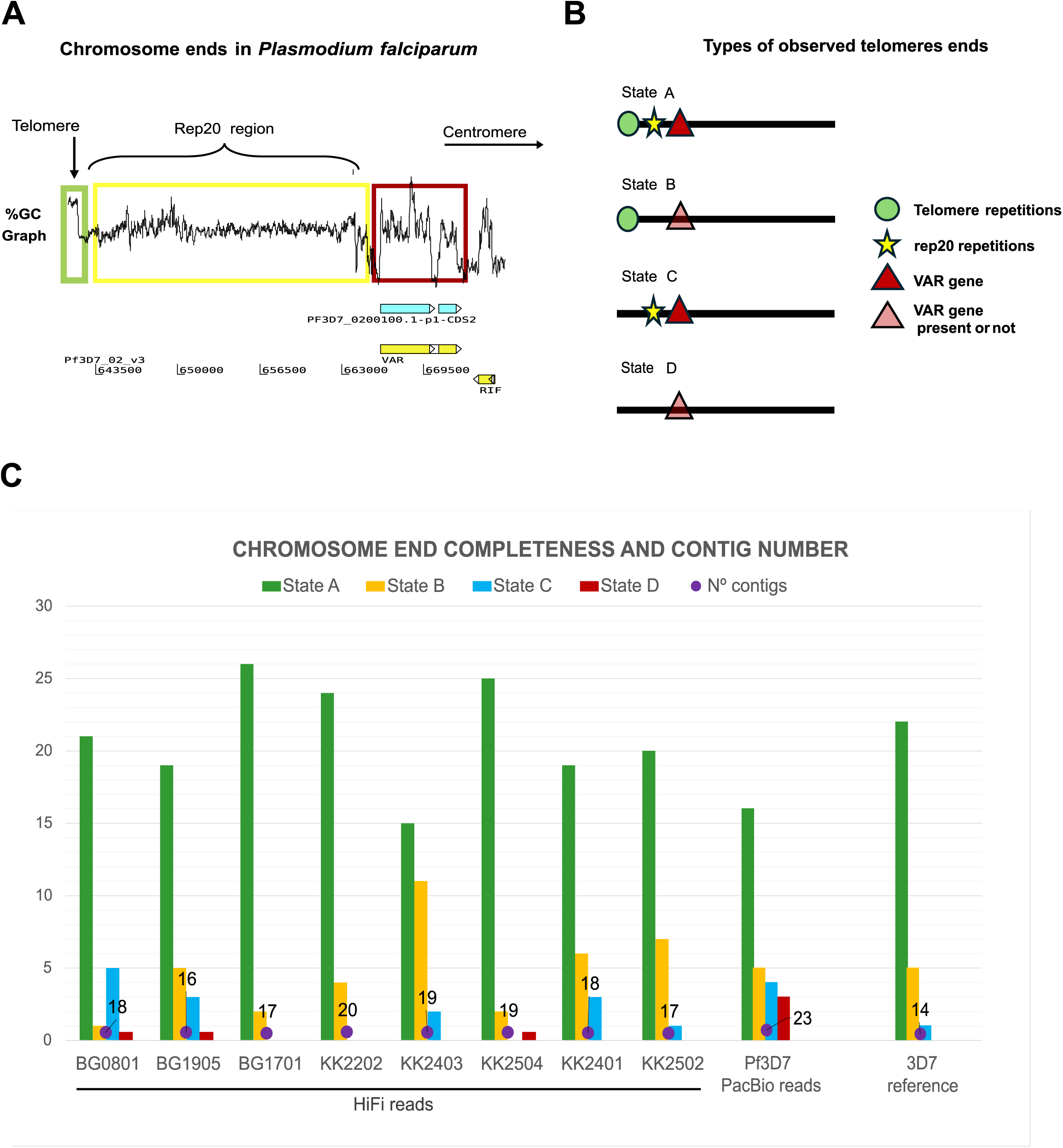
Defining complete chromosome-end completeness and assessing assembly quality in *P. falciparum* genomes generated with PacBio HiFi sequencing. **(A)** Schematic representation of a *P. falciparum* chromosome end. From the chromosome terminus inward, the structure comprises telomeric repeat sequences (green), the subtelomeric rep20 repeat region (yellow), and subtelomeric *var* genes (red). The genomic coordinates shown correspond to the Pf3D7 reference genome and illustrate the relative positions of these elements within a representative chromosome-end region. **(B)** Chromosome-end classification scheme used in this study. Chromosome ends were classified according to the presence or absence of telomeric repeats, the rep20 region and *var* genes. State A represents a complete chromosome end. State B represents chromosome ends containing telomeric repeats but lacking rep20 and containing or not a *var* gene. State C represents chromosome ends lacking telomeric repeats but retaining rep20 and a *var* gene. State D represents chromosome ends lacking both telomeric repeats and rep20 and containing or not a *var* gene. **(C)** Chromosome-end completeness and assembly quality metrics for the eight PacBio HiFi assemblies, the PacBio conventional 3D7 assembly, and the 3D7 reference genome. Stacked columns show the number of chromosome ends assigned to State A (green), State B (yellow), State C (blue), or State D (red). Purple circles indicate the total number of contigs in each assembly (left y-axis).

**Table 1.** Assembly statistics for the eight field-derived Plasmodium falciparum clones from Mali generated using PacBio HiFi sequencing. Genome size corresponds to the final assembly length after *ILRA* correction and Pilon polishing. N50 represents the minimum contig length for which contigs of that size or larger account for at least 50% of the total assembled genome. Gaps indicate unresolved regions remaining in the final assembly. The *P. falciparum* genome consists of 14 nuclear chromosomes, one mitochondrial genome, and one apicoplast genome (16 chromosomes total); therefore, contig numbers approaching 16 indicate highly contiguous assemblies.

| **Parasite clone** | **Size (Mbp)** | **Nº contigs** | **N50 (bp)** | **Gaps** |
|---|---|---|---|---|
| KK2504 | 23.59 | 19 | 1,668,498 | 0 |
| KK2202 | 23.60 | 20 | 1,643,388 | 0 |
| BG1701 | 23.51 | 17 | 1,673,768 | 0 |
| KK2403 | 23.15 | 19 | 1,671,314 | 0 |
| BG1905 | 23.42 | 16 | 1,684,515 | 0 |
| BG0801 | 23.67 | 18 | 1,656,947 | 1 |
| KK2401 | 23.31 | 18 | 1,687,130 | 0 |
| KK2502 | 23.42 | 17 | 1,693,086 | 2 |

We next compared these results with the PacBio-based 3D7 reference genome. Based on our classification, chromosome 14 was the only chromosome that remained incomplete (State D) in the reference assembly, whereas chromosome 14 was assembled to both chromosome ends in five of the eight de novo genomes (Fig. 1C). In addition, our analysis indicated that the Pf3D7 reference genome lacks rep20 sequences at the ends of chromosomes 5 and 14, corresponding to a State B classification for these chromosome ends (Fig. 1C).

To further investigate chromosome-end structure, we quantified telomeric repeat content across all chromosomal termini and visualized its distribution along each chromosome (Fig. 2A). Importantly, most chromosomal ends contain detectable telomeric repeats, indicating a high level of completeness across the assembled genomes. Consistent with the classification described above, the telomere-containing ends correspond to the combined counts of states A and B from Figure 1C, reinforcing the conclusion that most chromosome ends retain telomeric information. While most ends are characterized by moderate repeat content (30-100 repeats), regions with more extensive arrays (>100 repeats) are less frequent and vary among isolates. Overall, this pattern supports the robustness of the assemblies, and highlights the variability in the length of telomeric repeats among samples, rather than a general absence of telomeric sequences.

**Figure 2.**
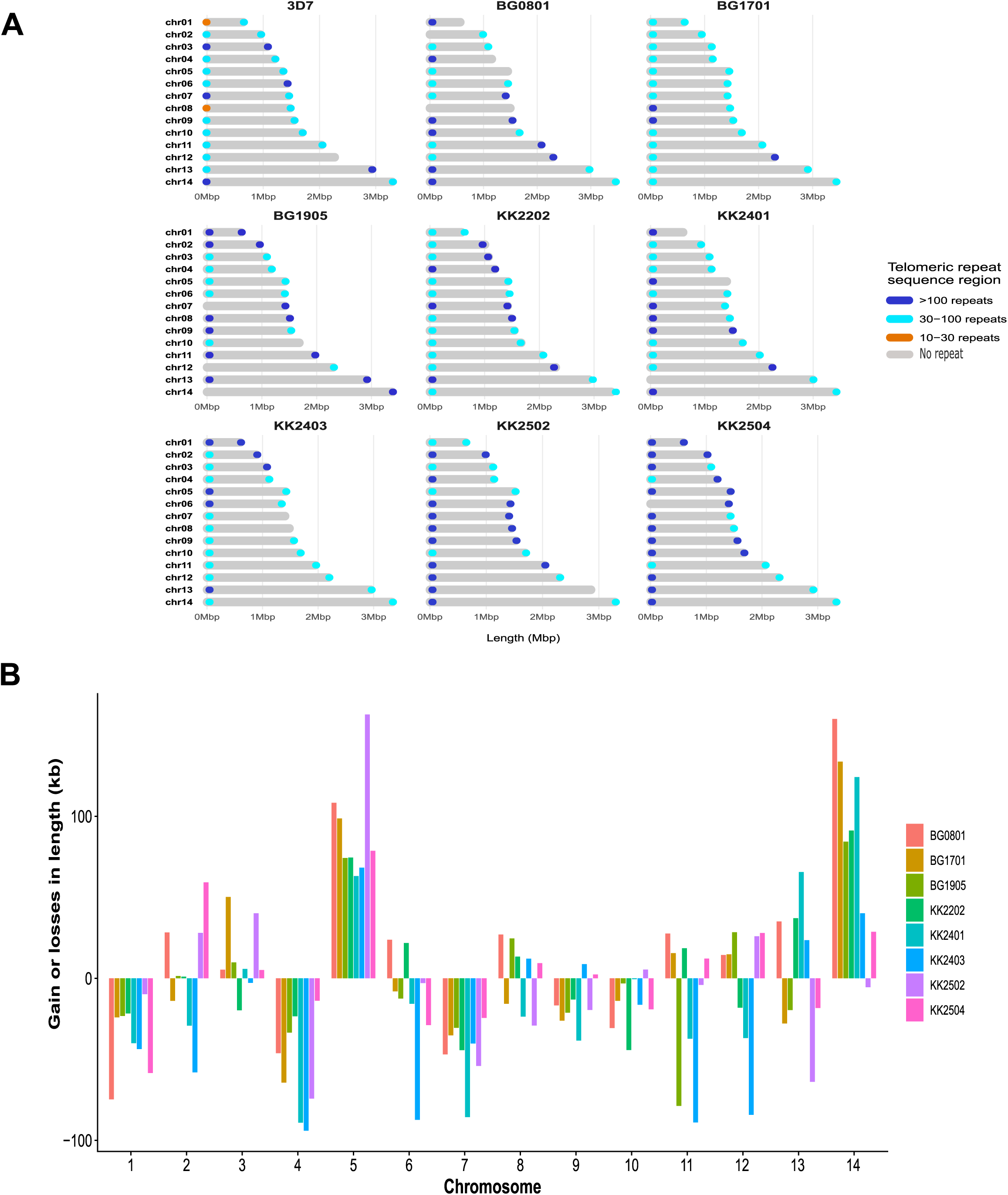
Telomere integrity and chromosome-length variation among *P. falciparum* isolates. **(A)** Distribution of telomeric repeat sequences across chromosome ends in the 3D7 reference genome and eight field-derived *P. falciparum* clones. Chromosomes (chr01-chr14) are shown according to their relative lengths, with telomeric repeat tracts represented by colored circles at chromosome termini. Green circles indicate >100 telomeric repeats, yellow circles indicate 30-100 repeats, and orange circles indicate 10-30 repeats. Chromosome ends without a symbol lacked detectable telomeric repeat sequences. This representation highlights variation in telomeric repeat abundance while preserving the relative organization of chromosomes within each genome. Telomeric repeat abundance is shown independently of the chromosome-end classification used in Fig. 1 and therefore does not necessarily indicate a complete chromosome end. **(B)** Chromosome-length differences relative to the 3D7 reference genome. gains or losses in chromosome length (kb) for each chromosome in individual assemblies. Positive values indicate chromosome expansion and negative values indicate contraction relative to the reference genome. Length variation is most pronounced in a subset of chromosomes, notably chromosomes 5 and 14.

Consistent with the variability observed at chromosome ends, we identified chromosome-specific differences in chromosome length across the assembled genomes (Fig. 2B). Most length changes were within approximately ±100 kb relative to the reference genome. Notably, chromosomes 5 and 14 exhibited recurrent length gains across multiple isolates and accounted for a substantial proportion of the chromosome-length variation observed in the dataset, whereas other chromosomes showed more heterogeneous patterns. To determine whether chromosome length variation was associated with broader structural rearrangements, we performed a synteny analysis against the 3D7 reference genome. Chromosome collinearity remained highly conserved across all samples (Fig. 3A). The circular synteny plots showed that gene order and overall chromosome organization were largely preserved throughout the core genome, with chromosome-wide synteny maintained between the assemblies and the 3D7 reference. Regions lacking alignment were predominantly restricted to subtelomeric domains, which are known to be highly variable and enriched in repetitive elements and antigenically important multigene families. These findings are consistent with the chromosome length differences described above and suggest that the observed gains and losses are driven primarily by expansions and contractions within subtelomeric regions rather than by large-scale structural alterations of the core genome. Collectively, the results indicate that although genome size varies among field isolates, the overall architecture of the conserved core genome remains largely intact, with most structural variability concentrated at chromosome termini.

**Figure 3.**
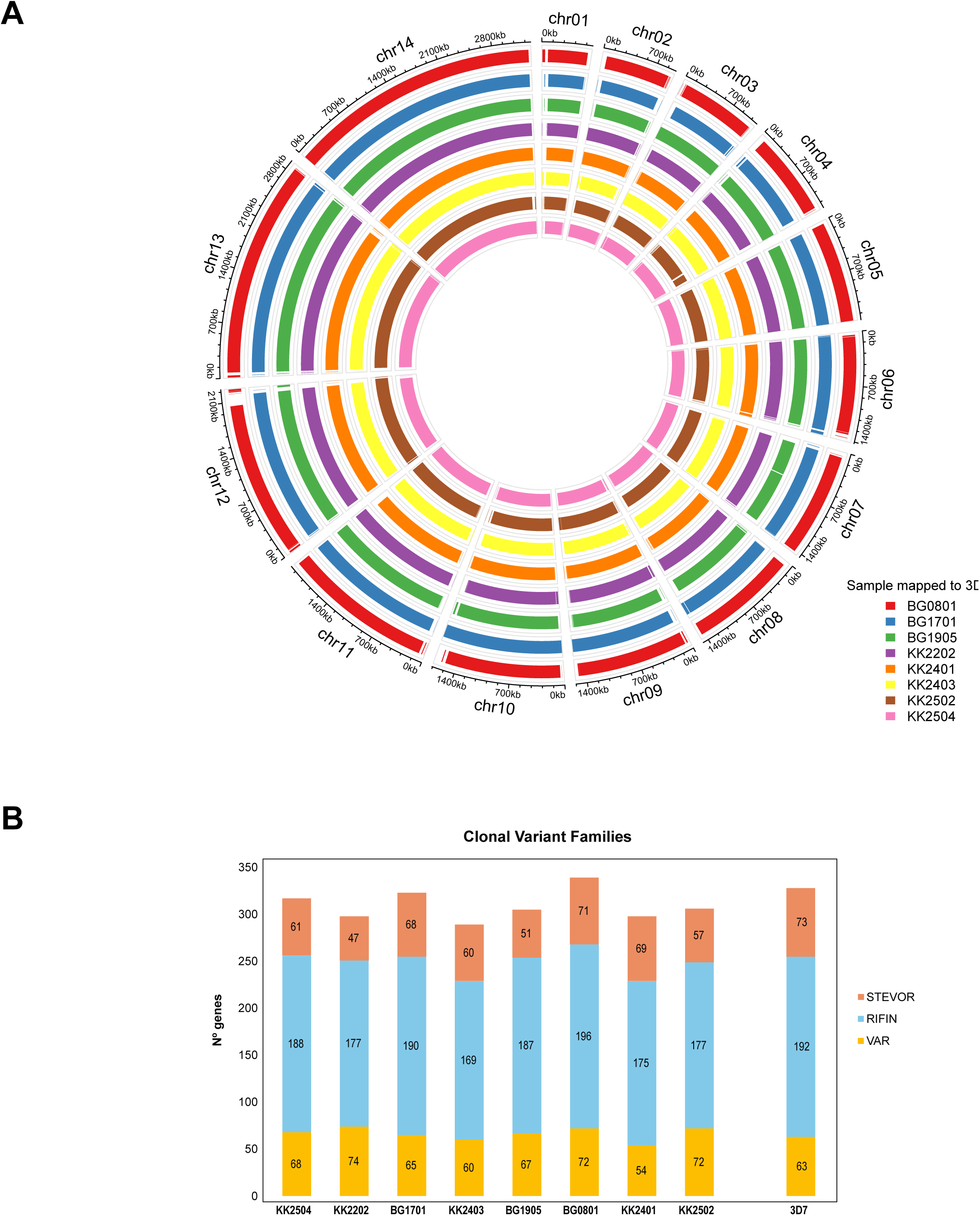
Genome organization and variation in major subtelomeric multigene families among *P. falciparum* clones. **(A)** *Circos* representation of the 14 *P. falciparum* chromosomes and whole-genome alignments of multiple isolates against the 3D7 reference genome. Each concentric ring corresponds to a different clone and illustrates the conservation of chromosomal structure as well as potential structural variations. Genome coordinates are shown in kilobases (kb). **(B)** Abundance of genes belonging to the major variant surface antigen families (*var*, *rifin*, and *stevor*) in each isolate. Stacked bars represent the contribution of each family to the total repertoire. Values within bars indicate the number of genes assigned to each family, enabling comparison among isolates and with the 3D7 reference genome.

Having established the structural completeness of the assemblies, we next evaluated annotation performance. Comparison of annotation strategies revealed substantial differences between *Liftoff* and *RATT*. Across the eight assemblies, *Liftoff* identified significantly more pseudogenes than *RATT* (mean 301 vs 166 pseudogenes per assembly; paired Wilcoxon signed-rank test, p = 0.0078, adjusted p = 0.047; rank-biserial correlation = -1.0). Conversely, *RATT* annotations showed significantly higher frameshift rates than *Liftoff* annotations (mean 3.39% vs 0.05%; paired Wilcoxon signed-rank test, p = 0.014, adjusted p = 0.047; rank-biserial correlation = 1.0). These findings demonstrated that the two annotation-transfer strategies produce consistently different annotation profiles, with *Liftoff* yielding more pseudogene calls and *RATT* generating a higher proportion of frameshifted gene models.

We next used hypervariable multigene families as an additional benchmark of assembly completeness in subtelomeric regions. We examined the number of annotated genes in multicopy and highly polymorphic gene families, such as *var*, *rifin*, and *stevor*. In all eight samples, the number of *var* genes ranges from 54 to 74, with a mean of 66. The mean number of *rifin* genes is 182, and for *stevor*, it is 60. The annotation with *RATT* performs better in detecting *var* genes. However, this higher sensitivity leads to an increase in the number of reading frame breaks. In contrast, although *Liftoff* appears to be less sensitive in subtelomeric regions, it demonstrates greater robustness across the core genome, which may be beneficial in applications that prioritize a low false-positive rate.

The newly assembled genomes contained *var, rifin*, and *stevor* gene repertoires comparable to those of the 3D7 reference genome (Figure 3B and Supplementary Table S2). Although the Malian assemblies tended to contain higher numbers of these multigene families than several other *P. falciparum* reference genomes (IT, SD01, GA01, KE01, Dd2, 7G8, KH01, and NF54), Wilcoxon rank-sum tests showed that these differences were not significant after correction for multiple testing (var: adjusted P = 0.12; *rifin*: adjusted P = 0.21; *stevor*: adjusted P = 0.12). Together, these results suggest that the multigene-family content of the Malian assemblies lies within the range of diversity represented by current reference genomes.

Taken together, PacBio HiFi long-read sequencing generated highly complete, high-quality genome assemblies for all samples analysed. Gene counts for *var*, *rifin*, and *stevor* genes varied among Malian assemblies, but numbers were comparable to those observed in the 3D7 and other reference genomes. These differences may reflect biological variation among parasite populations, differences arising during adaptation to in vitro culture, or a combination of both factors.

### 2. PacBio HiFi reads outperforms conventional PacBio and eliminates the need for short-read Illumina based polishing

Given the low error rate of PacBio HiFi reads, we propose that Illumina-based error correction becomes unnecessary. Consistent with this hypothesis, Figure 4A shows that the SNP error profile of HiFi reads is comparable to that of Illumina reads, which are commonly used to correct the higher SNP error rates associated with conventional PacBio sequencing. PacBio sequencing errors are particularly pronounced in repetitive sequence contexts, including homopolymer runs, which are a major source of indel errors in long-read data^23^ ^36^. To determine whether short-read correction can be omitted when using HiFi sequencing, we compared assemblies generated from PacBio HiFi and conventional PacBio reads at the same sequencing depth (200X). The assembly generated from conventional PacBio reads had a total size of 21.3 Mb, approximately 2.2 Mb shorter than the mean HiFi assembly size (23.5 Mb). Following assembly with *HiCanu*, the conventional PacBio dataset produced 305 contigs, compared with an average of 58 contigs for the HiFi datasets. After polishing with *Pilon* and correction with Illumina reads, the number of contigs was reduced to 20 for the conventional PacBio assembly and to an average of 18 for the HiFi assemblies (Supplementary Table S3).

**Figure 4.**
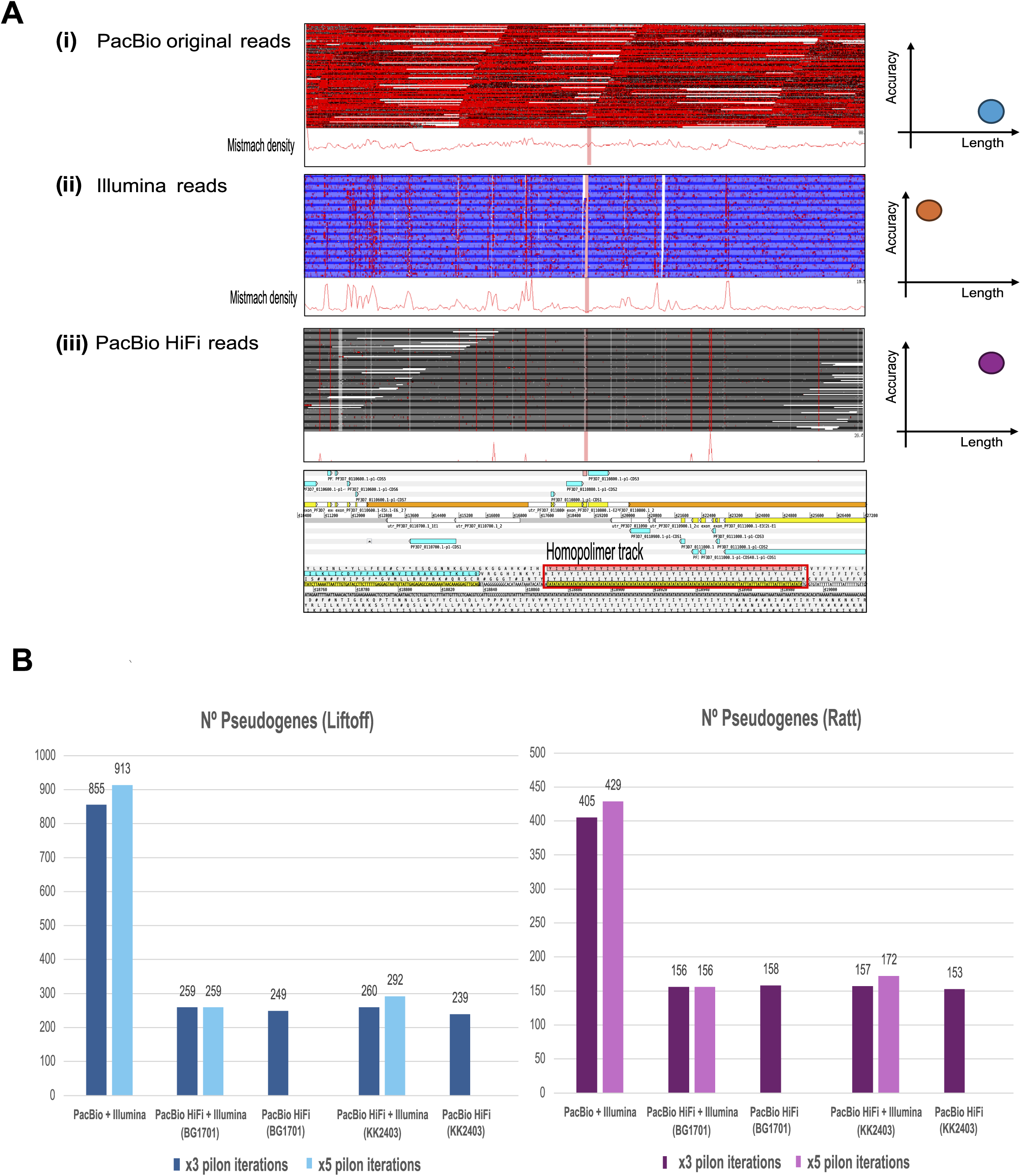
Assessing the impact of Illumina-based polishing on PacBio HiFi genome assemblies. **(A)** Comparison of sequencing error profiles in a representative *P. falciparum* genomic region containing a homopolymer tract (red box). Read alignments, consensus sequences (pink bars), and mismatch-density tracks (red histograms) are shown for (i) conventional PacBio reads, (ii) Illumina reads, and (iii) PacBio HiFi reads. Conventional PacBio reads display a higher frequency of sequence discrepancies, particularly around homopolymer regions, whereas PacBio HiFi reads exhibit an error profile comparable to that of Illumina reads. **(B)** Effect of polishing and correction on assembly annotation quality. The number of pseudogenes and the proportion of genes containing frameshifts were assessed using the *Liftoff* and *RATT* annotation pipelines before and after *ILRA* correction and *Pilon* polishing. Dark and light bars represent results after three and five Pilon polishing iterations, respectively. All assemblies were generated using 200X sequencing coverage to enable direct comparison. Conventional PacBio assemblies contained substantially more pseudogenes and frameshifts than PacBio HiFi assemblies, whereas Illumina-based correction and additional *Pilon* iterations produced only minor changes in HiFi assemblies (see Supplementary Table S3).

Assemblies generated from HiFi reads also contained fewer annotated pseudogenes than those generated from the conventional PacBio reads (Figure 4B; Supplementary Table S3). Using the *Liftoff* annotation pipeline, the corrected conventional PacBio assembly contained 855 pseudogenes, compared with 259 and 260 for the two HiFi assemblies. Using *RATT*, the corresponding values were 405 and 156-157 pseudogenes, respectively. Similarly, the conventional PacBio assembly exhibited a higher proportion of genes containing frameshifts. With *Liftoff* annotation, frameshifts were detected in approximately 1.1% of annotated genes in the conventional PacBio assembly, compared with approximately 0.04% in the HiFi assemblies. Using *RATT*, the corresponding values were approximately 5.0% and 3.0%, respectively (Supplementary Table S3).

To directly assess the contribution of Illumina-based correction, we compared assembly statistics before and after *ILRA* correction for two representative HiFi genomes (BG1701 and KK2403) and contrasted these results with assemblies generated from conventional PacBio reads (Supplementary Table S3; Supplementary Figure S1B). Illumina-based correction produced only minor changes in assembly metrics. Consistent with this observation, paired comparisons detected no significant differences in pseudogene counts using either *Liftoff* (P = 0.50) or *RATT* (P = 1.0), while *RATT*-derived frameshift frequencies also remained unchanged (P = 1.0). Liftoff-derived frameshift estimates were identical before and after correction and could therefore not be statistically evaluated. These findings indicated that Illumina correction provided no measurable improvement to PacBio HiFi assemblies.

Increasing the number of Pilon polishing iterations from three to five rounds did not significantly alter pseudogene counts or frameshift frequencies under either annotation strategy (all Wilcoxon signed-rank tests, P = 1.0). These findings indicated that additional polishing beyond three iterations provided no measurable improvement in assembly annotation quality.

Together, these results indicate that Illumina-based correction provides little additional benefit for PacBio HiFi assemblies, whereas it remains important for assemblies generated from conventional PacBio reads. HiFi sequencing therefore enables the generation of highly contiguous and accurate *P. falciparum* genome assemblies while substantially reducing the dependence on complementary short-read sequencing.

### 3. PacBio HiFi reads reduce the coverage requirements 4-fold for the reconstruction of complete *P. falciparum* genome assemblies

Optimal sequencing depth varies with the study design, organism, and sample characteristics. Identifying the required depth in advance can reduce costs and analysis effort. We therefore evaluated the coverage required for PacBio HiFi reads to generate high-quality assemblies. We tested different sequencing depths following the Lander-Waterman equation. Figure S2A shows the theoretical coverage required for our genome, based on the average read length, to achieve an assembly with approximately zero gaps. According to this estimation, a coverage of ∼22X would be sufficient to obtain a gapless genome. To explore the effect of coverage on assembly quality, we deliberately generated data subsets at lower and higher depths than this theoretical value, creating seven coverage levels: 5X, 10X, 25X, 50X, 100X, and 200X. We used the same parameters described in the previous section to evaluate these assemblies.

Looking at genome size, *HiCanu* failed to produce a complete *P. falciparum* assembly at the two lowest coverages tested (5X and 10X). In contrast, assembly contiguity, as measured by N50, remained comparable to that achieved at the original 200X coverage across all tested coverages and only declined substantially below 20X coverage (Figure S2B; Supplementary Table S4). Following assembly with *HiCanu*, the number of contigs increased as coverage decreased, but remained relatively stable at an average of 23 contigs down to 25X coverage (Supplementary Table S4). Subsequently, in the second stage of the pipeline, after applying *ILRA* and *Pilon*, the number of contigs stabilized at an average of 18 for all samples and coverages. This result is consistent with tests performed on the PacBio original dataset comparing 450X and 200X coverages, suggesting an improved assembly continuity and effective consolidation of contigs throughout the genomic assembly process.

Sequencing coverage also affected assembly contiguity prior to ILRA correction. Lower sequencing coverage generally resulted in more fragmented HiCanu assemblies, as reflected by an increased number of pre-corrected contigs. This effect was significant (χ² = 16.40, P = 9.4 × 10⁻□, Kendall’s W = 0.91), indicating a strong dependence of initial assembly contiguity on sequencing depth. However, after ILRA correction, contig numbers did not differ significantly among coverage levels (χ² = 6.25, P = 0.10), showing that *ILRA* largely compensated for the fragmentation present in lower-coverage assemblies (Supplementary Table S5).

The occurrence of assembly gaps appeared independent of sequencing coverage (Supplementary Table S4). Most assemblies contained no gaps, and the few gaps observed were limited to a single gap per assembly, ranging from 10 to 76 bp in length. The low frequency and small size of these gaps indicate that reducing sequencing coverage has little impact on overall assembly quality.

We next evaluated the impact of sequencing depth on chromosome-end completeness. Analysis of chromosome-end structure revealed a progressive decline in fully resolved chromosome ends (State A) as sequencing coverage decreased, accompanied by a corresponding increase in chromosome ends lacking telomeric repeats (State C) (Figure S2C). Specifically, the average proportion of complete chromosome ends declined from 77.2% at 200X coverage to 36.4% at 25X coverage. In parallel, the proportion of chromosome ends classified as complete but lacking telomeric repeats increased from 5.9% to 45.8% over the same coverage range. A similar pattern was observed for incomplete chromosome ends: the proportion of incomplete ends containing telomeric repeats decreased from 14.9% at 200X coverage to 7.1% at 25X coverage, while the proportion lacking telomeric repeats increased from 1.8% to 10.1%. Friedman tests confirmed a significant effect of coverage on both State A (χ² = 16.97, P = 7.2 × 10⁻□, Kendall’s W = 0.94) and State C chromosome ends (χ² = 17.53, P = 5.5 × 10⁻□, Kendall’s W = 0.97), indicating a strong and consistent reduction in chromosome-end completeness at lower sequencing depths. In contrast, the proportions of State B and State D chromosome ends did not differ significantly across coverage levels (Supplementary Table S5). Together, these results indicated that reduced sequencing depth primarily compromised chromosome-end completeness and telomere recovery.

The effect of sequencing coverage on annotation quality was evaluated using pseudogene counts and frameshift frequencies (Supplementary Table S4). Although pseudogene counts varied among isolates and between annotation methods, no significant effect of coverage was detected for either *Liftoff* (χ² = 4.20, P = 0.241) or *RATT* annotations (χ² = 4.50, P = 0.212). Frameshift frequencies remained highly consistent across coverage levels, averaging approximately 0.04% with Liftoff and 3.1% with *RATT*. Coverage did not significantly affect *RATT*-derived frameshift frequencies (χ² = 6.60, P = 0.086). A significant effect was detected for *Liftoff*-derived frameshift frequencies (χ² = 8.57, P = 0.036; Kendall’s W = 0.48), although the moderate effect size and absence of a consistent directional trend suggest limited biological relevance. Overall, these findings indicate that reduced sequencing coverage had little impact on gene annotation quality despite its effects on chromosome-end completeness and initial assembly contiguity.

We applied the same coverage reductions to the 3D7 sample obtained with non-HiFi PacBio reads. In this case, only the 200X coverage was able to generate results of similar quality. For the other coverages (100X, 50X, and 25X), *HiCanu* failed to assemble the genome. This is likely due to the high per-read error rate of the conventional PacBio reads, which requires higher coverage to provide enough overlapping information for accurate assembly. By comparing the 200X sample statistics of these reads with the HiFi reads of the same coverage, we found a similar number of contigs after *ILRA* (19 in PacBio HiFi vs. 23 in PacBio original). The number of gaps was 301, with sizes of up to 5,403 bp. Regarding the completeness of chromosome ends, 71.4% of the chromosome ends were in state A (∼ complete), 21.4% in state B, and 3.5% in states C and D (Supplementary Table S4). Regarding the two annotation methods, we observe a clear increase both in the number of pseudogenes (943 in *Liftoff* versus 462 in *RATT*) and in the percentage of frameshifts, which stands at 1.4% for the Liftoff annotation and 6.16% for the *RATT*-generated annotation (Supplementary Table S4). These results are in line with the trend observed with the higher coverage (200X), where these parameters are significantly higher when using the original reads. Therefore, the use of the original reads does not match the quality of results obtained with the HiFi reads even at the standard 200X coverage level defined by *HiCanu*.

This analysis also demonstrates remarkable consistency in the recovery of multigene families, regardless of the coverage level used. The results indicate that the number of *var* genes (identified using the LARSFADIG motif), remains similar across a coverage range from 200X to 25X (Table 2), as also observed for the *rifin* and *stevor* families (Supplementary Table S4C). For example, for BG0801, the count of LARS motifs stabilized at 69 copies for all coverages above 50X, showing negligible variations (±1 copy) when decreasing to 25X in most samples (Table S4C). This stability suggests that the architecture of the hypervariable regions is robustly captured even under conditions of lower depth.

**Table 2.** Effect of sequencing depth on *var* gene recovery in PacBio HiFi assemblies. . The number of var genes was determined using the LARSFADIG motif across assemblies generated from subsampled datasets at four sequencing depths (25X, 50X, 100X, and 200X coverage). Gene counts remained largely stable across coverage levels, indicating robust recovery of hypervariable subtelomeric gene families even at reduced sequencing depth.

| **Sample** | **Coverage** | **Nº VAR** |
|---|---|---|
| KK2504 | 200X | 69 |
|  | 100X | 69 |
|  | 50X | 69 |
|  | 25X | 68 |
| KK2202 | 200X | 74 |
|  | 100X | 71 |
|  | 50X | 71 |
|  | 25X | 71 |
| BG1701 | 200X | 65 |
|  | 100X | 65 |
|  | 50X | 66 |
|  | 25X | 66 |
| KK2403 | 200X | 60 |
|  | 100X | 59 |
|  | 50X | 58 |
|  | 25X | 59 |
| BG1905 | 200X | 67 |
|  | 100X | 68 |
|  | 50X | 69 |
|  | 25X | 69 |
| BG0801 | 200X | 72 |
|  | 100X | 72 |
|  | 50X | 72 |
|  | 25X | 72 |

Collectively, these tests demonstrate that it is possible reduce the sequencing depth by 4-fold of the previously established threshold. This substantial decrease in sequencing requirements has key implications for improving efficiency and reducing computational and operational costs in genome assembly projects.

### 4. Detection of drug resistance-associated variants in Malian parasite genomes

To evaluate the utility of the assemblies for genomic surveillance, we screened all eight genomes for known drug resistance-associated markers (Supplementary Table S6). Several validated resistance mutations were readily identified in the de novo assemblies, demonstrating accurate reconstruction of clinically relevant loci. Notably, mutations associated with sulfadoxine-pyrimethamine resistance were common, including the *pfdhfr* mutations N51I, C59R, and S108N, and the *pfdhps* mutations A437G and S436A, although their distribution varied among parasite clones. The chloroquine resistance-associated *pfaat1* S258L mutation was also detected in most genomes, while the *pfmdr1*Y184F variant was present in five clones. In contrast, no validated artemisinin-resistance mutations were detected in *pfk13*, and the major chloroquine-resistance mutation *pfcrt* K76T was absent from all assemblies. Several candidate or investigational resistance markers were also identified, including variants in *pfubp1*, *pfpi4k*, *pfap2-mu*, and *pfmdr2*. Varation in quinine-resistance-associated pfnhe1 microsatellite motifs was limited among the Malian assemblies, with two major repeat profiles observed across the isolates (Supplementary Table S7). These results show that PacBio HiFi assemblies enable reliable recovery of both validated and emerging drug-resistance markers directly from de novo genome assemblies.

## DISCUSSION

PacBio HiFi sequencing, combined with an optimized assembly workflow, enabled the generation of highly contiguous, near telomere-to-telomere assemblies from field-derived *Plasmodium falciparum* clones while reducing sequencing, computational, and analytical requirements. The assemblies had a mean size of 23.5 Mb, closely matching the 23.6-Mb 3D7 reference genome, and were highly contiguous, with a small number of contigs. Together, these features indicate minimal sequence loss and high structural integrity. Beyond improving assembly performance, the resulting genomes provided a detailed view of chromosome-end architecture, revealing extensive variation in telomeric repeat content and subtelomeric organization among clones of parasites isolated recently from the field. The assemblies also enabled robust characterization of hypervariable multigene families, demonstrating substantial diversity in *var*, *rifin*, and *stevor* repertoires while preserving the overall structural conservation of the core genome.

### Assembly performance and workflow optimization

Recovery of chromosome ends and subtelomeric regions is a key measure of genome assembly completeness in *P. falciparum*, because these regions have historically been difficult to assemble owing to their highly repetitive organization, extreme AT richness, and enrichment in structurally-variable multigene families involved in antigenic variation^11^ ^37^ . In this study, approximately 74% of chromosome ends were recovered as complete telomere-containing sequences, with some parasite genomes exceeding 90% completeness. The widespread detection of telomeric repeats across chromosome termini therefore shows that HiFi sequencing, combined with our assembly pipeline, resolves genomic regions that have remained fragmented in previous assemblies.

This chromosome-end resolution was accompanied by consistently high genome continuity across samples. Compared with recently reported HiFi-based assemblies^13^, our assemblies showed comparable or improved continuity and contained a mean of only 18 contigs; in several cases, only one or two gaps remained across the entire genome (compared to the reference 3D7 genome). These results indicate that high-fidelity long reads can generate near-complete *P. falciparum* genomes from field parasites without extensive post-processing of the sequence data.

The accuracy of HiFi reads also removed the requirement for Illumina-based polishing that accompanied earlier long-read technologies^38^. When the same workflow and optimization strategy were applied, assemblies generated from conventional PacBio reads were more fragmented and yielded more pseudogenes and annotation errors than those generated from HiFi reads. Additional short-read polishing produced little measurable improvement when HiFi reads were available, indicating that HiFi sequencing alone is sufficient to generate high-quality genome assemblies. This substantially simplifies the workflow while reducing sequencing and computational costs.

Coverage analysis further demonstrated the efficiency of the approach. Assemblies generated at 25-30X coverage were statistically indistinguishable from those obtained at higher sequencing depths for most assembly and annotation metrics, including pseudogene counts and frameshift frequencies. The only assembly feature consistently affected by reduced coverage was chromosome-end completeness, where significant decreases in complete chromosome ends and increases in telomere-deficient chromosome ends were observed. Thus, 25-30X coverage appears sufficient for most genomic applications, whereas deeper sequencing is advantageous when complete reconstruction of chromosome termini is required. Consequently, coverage requirements should be adapted to the biological question, with higher depth providing the greatest benefit for studies focused on telomere architecture and chromosome-end dynamics.

### Subtelomeric plasticity and chromosome-end dynamics

Our data highlight the extensive structural variability present at the chromosome ends of *P. falciparum*. Although chromosome synteny was highly conserved across the core genome, homologous chromosomes differed in length by up to approximately 100 kb among isolates. Notably, recurrent length increases were observed for chromosomes 5 and 14 across multiple genomes and were primarily associated with differences in subtelomeric regions rather than the conserved chromosome core. These observations are consistent with previous studies showing that structural variation in *P. falciparum* is concentrated at chromosome ends and that subtelomeric domains represent the most variable regions of the genome ^5,37,39,40^.

Variation in telomeric repeat length was also common among parasite genomes and is likely to reflect biological differences rather than assembly artefacts. Telomere length in *P. falciparum* is influenced by telomerase activity, recombination, telomere healing, and chromosome-end repair processes^41,42^ . The widespread recovery of telomeric repeats in our assemblies, together with the substantial variation observed in subtelomeric size, further supports the view that *P. falciparum* chromosome ends are highly variable genomic compartments.

This plasticity is further reinforced by repetitive elements such as rep20, which facilitate ectopic homologous recombination and contribute to subtelomeric rearrangements^40,43^ . Through recombination, duplication, expansion, and contraction, chromosome ends act as evolutionary hotspots that generate genomic diversity while preserving the stability of the conserved core genome^39,40^. The observed size differences and structural variability are therefore consistent with natural patterns of chromosome-end evolution in field isolates. The distinct chromosome-end profile observed in the genome of parasite clone KK2403 further illustrates the diversity present among parasites recently isolated from the field. Although this parasite genome contained an elevated proportion of chromosome ends lacking rep20, inspection of the underlying HiFi reads supported these structures, suggesting that they may represent genuine biological variation rather than assembly artefacts.

More recently isolated parasites may also preserve a broader spectrum of subtelomeric diversity than laboratory-adapted parasites that have been maintained *in vitro* for long periods of time. Prolonged *in vitro* culture has been associated with loss of subtelomeric sequences and virulence-associated genes, potentially underestimating the complexity present in natural populations^44^ . Accordingly, the extensive variation among our parasite genomes likely reflects biologically relevant diversity maintained under natural transmission and immune selection^5,39^.

*Accurate reconstruction of hypervariable and clinically relevant genomic loci*A central advantage of highly contiguous assemblies is the improved characterization of hypervariable gene families involved in antigenic variation and immune evasion. The *var*, *rifin*, and *stevor* families are concentrated in subtelomeric regions and are among the most challenging genomic features to assemble accurately^5,39^. Our assemblies recovered numbers of *var* gene comparable to those reported in high-quality reference genomes and recent HiFi studies^11,13^, while providing improved structural resolution of these loci. Because most chromosome ends were assembled completely, internal and subtelomeric *var* genes could be distinguished with high confidence, a distinction often blurred in fragmented assemblies^11,39^. Recovery of *rifin* and *stevor* genes was also consistently high, with several assemblies containing more copies than in previously reported parasite genomes. This likely reflects the improved assembly contiguity achieved in subtelomeric and repetitive regions, reducing the collapse of closely related gene copies.

These assemblies also provide a detailed view of natural variation in chromosome-end architecture, telomeric repeat content, and multigene family repertoires among recently field-adapted parasites. Because subtelomeric multigene families are major determinants of parasite virulence, immune evasion, and host-parasite interactions, their accurate reconstruction is important for studies of parasite evolution, population diversity, and transmission biology.

Beyond resolving these complex genomic regions, the assemblies proved suitable for the characterization of antimalarial drug-resistance loci. Multiple validated mutations associated with sulfadoxine-pyrimethamine resistance were identified across the Malian parasite genomes. Whereas validated artemisinin-resistance mutations in *pfk13* were not detected, we identified several candidate and background artemisinins-related mutations. The accurate recovery of both established and candidate resistance-associated variants highlights the utility of PacBio HiFi assemblies for genomic surveillance. As genome sequencing becomes increasingly integrated into malaria surveillance programs, highly contiguous field-derived assemblies offer the opportunity to simultaneously characterize structural genome variation and clinically relevant resistance markers within contemporary parasite populations.

## CONCLUSION

PacBio HiFi sequencing, combined with an optimized assembly workflow, provides a practical and scalable strategy for generating near telomere-to-telomere *P. falciparum* genome assemblies. The ability to obtain highly contiguous assemblies without extensive polishing, together with substantially reduced sequencing requirements, lowers the technical and economic barriers to the routine production of high-quality malaria parasite reference genomes.

These advances facilitate the incorporation of geographically-diverse field parasites into genomic studies and reference genome collections. By improving access to complete chromosome-end and subtelomeric assemblies, this approach provides a stronger foundation for investigations of parasite evolution, antigenic variation, genome plasticity, and population diversity, while supporting genomic surveillance efforts in malaria-endemic regions.

## Supplementary Figure Legends

**Fig. S1. PacBio HiFi read-length distribution and chromosome-end completeness across sequencing strategies.**

**(A)** Distribution of PacBio HiFi read lengths for eight field-derived P. falciparum clones. Violin plots show the density and variability of read lengths obtained for each clone, with embedded boxplots indicating the median and interquartile range. The total number of reads analysed for each clone is indicated above the corresponding violin plot.

**(B)** Comparison of chromosome-end completeness and assembly quality metrics among assemblies generated using PacBio HiFi reads alone, PacBio HiFi reads with Illumina-based correction, conventional PacBio reads with Illumina-based correction, and the 3D7 reference genome. Columns show the number of chromosome ends assigned to State A (telomere+/rep20+/*var*+; green), State B (telomere+/rep20−/*var*+−; yellow), State C (telomere−/rep20+/*var*+; blue), or State D (telomere−/rep20−/ *var*+−; red). Purple and dark red circles indicate the total number of contigs and pseudogenes identified in each assembly. Left and right y-axes correspond to number of contigs and number of pseudogenes, respectively.

**Fig. S2. Effect of sequencing coverage on *P. falciparum* genome assembly quality using PacBio HiFi reads.**

**(A)** Expected number of assembly gaps as a function of sequencing coverage, estimated using the Lander-Waterman model. The curve shows the decline in expected gaps with increasing coverage (50X, 100X, 150X, and 200X). The blue point indicates the minimum coverage at which the expected number of gaps reaches zero.

**(B)** N50 values obtained at different sequencing coverages (10X, 20X, 50X, 100X, and 200X) using different sequencing coverage levels. Colored lines represent different assembly methods, and error bars indicate variability among replicates.

**(C)** Completeness of chromosome ends in assemblies generated at different sequencing coverages (25X, 50X, 100X, and 200X). Stacked bars show the proportion of chromosome ends classified as complete (State A, green), lacking REP20rep20 (State B, yellow), lacking telomeric repeats (State C, blue), or containing only a *var* gene (State D, red). Reduced coverage results in fewer complete chromosome ends and a higher proportion of telomere-deficient ends.

## SUPPLEMENTARY TABLE LEGENDS

**Table S1. Summary of PacBio long-read sequencing data from clonal and isolate-derived *P. falciparum* samples.**

Summary of sequencing output generated from clonal and synthetic *P. falciparum* samples using PacBio long-read sequencing. Reported metrics include sample identifier, biological source, barcode assignment, number of <u>polymerase</u> reads, total base pairs generated, mean read length, average barcode quality, and estimated genome coverage.

**Table S2. Assembly statistics, annotation metrics, and chromosome-end completeness of *P. falciparum* isolates compared with the 3D7 reference genome.**

**(A)** Assembly statistics following initial genome assembly with *HiCanu*.

**(B)** Assembly statistics and annotation metrics after *Pilon* polishing. Reported values include contig number, mean contig size, largest contig length, N50, gap content, gene annotation statistics obtained with *Liftoff* and *RATT*, pseudogene counts, frameshift frequencies, and numbers of genes belonging to the *var*, *rifin*, and *stevor* multigene families.

**(C)** Chromosome-end completeness across isolates relative to the 3D7 reference genome. Chromosome termini are classified as complete (State A), lacking rep20 (State B), lacking telomeric repeats (State C), or containing only a *var* gene (State D). Summary statistics and proportions are provided for each isolate.

**Table S3. Assembly quality and genome annotation following Illumina-based polishing.**

**(A)** Assembly and annotation metrics for *P. falciparum* genomes generated from PacBio HiFi and conventional PacBio data after Illumina-based correction. Assembly statistics include total assembly length, contig number, mean contig size, largest contig, N50-N100 values, number of ambiguous base pairs (N count), and gap counts. Annotation metrics include gene counts, pseudogene counts, coding and non-coding genes, genes with assigned functions, frameshift frequencies, and representation of the *var*, *rifin*, and *stevor* multigene families.

**(B)** Comparison of assembly and chromosome-end status after three and five rounds of Pilon polishing. Chromosome ends are categorized as complete (State A), lacking rep20 (State B), lacking telomeric repeats (State C), or containing only a *var* gene (State D), enabling evaluation of the impact of iterative polishing.

**Table S4. Influence of sequencing coverage on assembly quality, genome annotation, and multigene family representation.**

**(A)** Assembly statistics generated at different sequencing coverages following assembly with *HiCanu* and correction with *ILRA*. Metrics include contig number, total assembly size, N50, gap count, and number of ambiguous base pairs.

**(B)** Classification of chromosome ends according to four structural states: complete (State A), lacking rep20 (State B), lacking telomeric repeats (State C), and containing only a *var* gene (State D). Pseudogene numbers obtained using the default annotation pipeline and RATT are also reported.

**(C)** Annotation quality metrics, including frameshift frequencies and identification of parasite-specific multigene families (*var*, *rifin*, and *stevor*). The number of genes, detected motifs, and contigs containing each family are reported across sequencing coverages and samples, allowing evaluation of assembly completeness and annotation accuracy as a function of sequencing depth.

**Table S5. Statistical evaluation of assembly construction, polishing, annotation transfer methods, sequencing coverage, and multigene family content.**

Paired comparisons were evaluated using two-sided Wilcoxon signed-rank tests. Coverage-dependent effects were assessed using Friedman tests across assembly coverages (25X, 50X, 100X and 200X). Comparisons between Malian assemblies and PlasmoDB reference genomes were performed using Wilcoxon rank-sum tests. Effect sizes are reported as rank-biserial correlation (RBC) or Kendall’s coefficient of concordance (W). Adjusted p-values correspond to Benjamini-Hochberg correction where applied. Statistically significant results (p < 0.05) supported significant effects of sequencing coverage on chromosome-end completeness (States A and C), contig numbers before *ILRA* processing, and *Liftoff*-derived frameshift rates, as well as systematic differences between Liftoff and *RATT* annotation outcomes.

**Table S6. Antimalarial drug resistance variants.** Amino acid variants detected in the assembled Plasmodium falciparum genomes at loci associated with resistance or altered susceptibility to approved and investigational antimalarial compounds. For each marker, the corresponding gene, drug association, resistance classification, reference and observed amino acids, and variant status are reported for each isolate.

**Table S7. Microsatellite polymorphisms.** Repeat numbers of the quinine-resistance-associated microsatellite motifs within the *pfnhe1* locus for each assembled isolate. Drug association and resistance classification are indicated for each motif.

## DATA AVAILABILITY

### Underlying data

All sequencing data, assemblies, and associated metadata have been deposited in public repositories to facilitate reproducibility and reuse.

Illumina whole-genome sequencing data and PacBio HiFi sequencing data generated for the eight *P. falciparum* clones have been deposited in ENA under accession PRJEB98187.

### Extended data

Supplementary tables and figures supporting the findings of this work, including assembly statistics, annotation metrics, chromosome-end classifications, and coverage-reduction analyses, are available as part of the article’s supplementary material.

## SOFWARE AVAILABILITY

Custom scripts used for the analyses described in this work, including pseudogene/frameshift assessment and multigene family identification, are publicly available from GitHub: https://github.com/trn30/Assemblies_Mali_field_genotypes. All third-party software used in the study is freely available and cited in the Methods section. Software versions and parameters are reported throughout the manuscript.

## AUTHOR CONTRIBUTIONS

Conceptualization: E.G.-D. and T.D.O.; Methodology: B.D.-T., E.G.-D. and T.D.O.; Software: B.D.-T., M.C.M-C and T.D.O.; Formal analysis: B.D.-T., M.C.M-C and T.D.O.; Laboratory Experiments: B.D.-T., S.P., and L.C.R.-C.; Resources: M.D.; Data curation: B.D.-T.; Visualization: B.D.-T.; Supervision: T.D.O. and E.G.-D.; Funding acquisition: E.G.-D., M.D., L.C.R.-C, T.D.O. and E.G.-D.; Writing – original draft: B.D.-T. and E.G.-D.; Writing – review and editing: all authors. All authors read and approved the final manuscript.

## COMPETING INTERESTS

The authors declare that they have no competing interests.

## GRANT INFORMATION

This work was supported by the Wellcome Trust ([104111/Z/14/Z&A, 225254/Z/22/Z]) and by the Spanish Ministry of Science and Innovation [grant no. PID2019-111109RB-I00] and by La Caixa Foundation—Health Research Program [grant no. HR20-00635]. B.D.-T. was supported by [grant no. PRE2020-095076].

The funders had no role in study design, data collection and analysis, decision to publish, or preparation of the manuscript.

## ACKNOWLEDGMENTS

We thank the study participants and clinical teams involved in sample collection in Mali. We are grateful to members of the participating laboratories for technical assistance, parasite culture support, and helpful discussions. Computational analyses were performed using resources provided by IPBLN-CSIC. This manuscript is part of the doctoral thesis of Bárbara Díaz Terenti in the Ph.D. program of molecular biology and biochemistry of the University of Granada.

## Notes

### Competing Interest Statement

The authors have declared no competing interest.

