## Supplementary Figure 1 for "Improved long-read sequencing technology reveals full genome architecture of Malian *Plasmodium falciparum* field isolates"

A

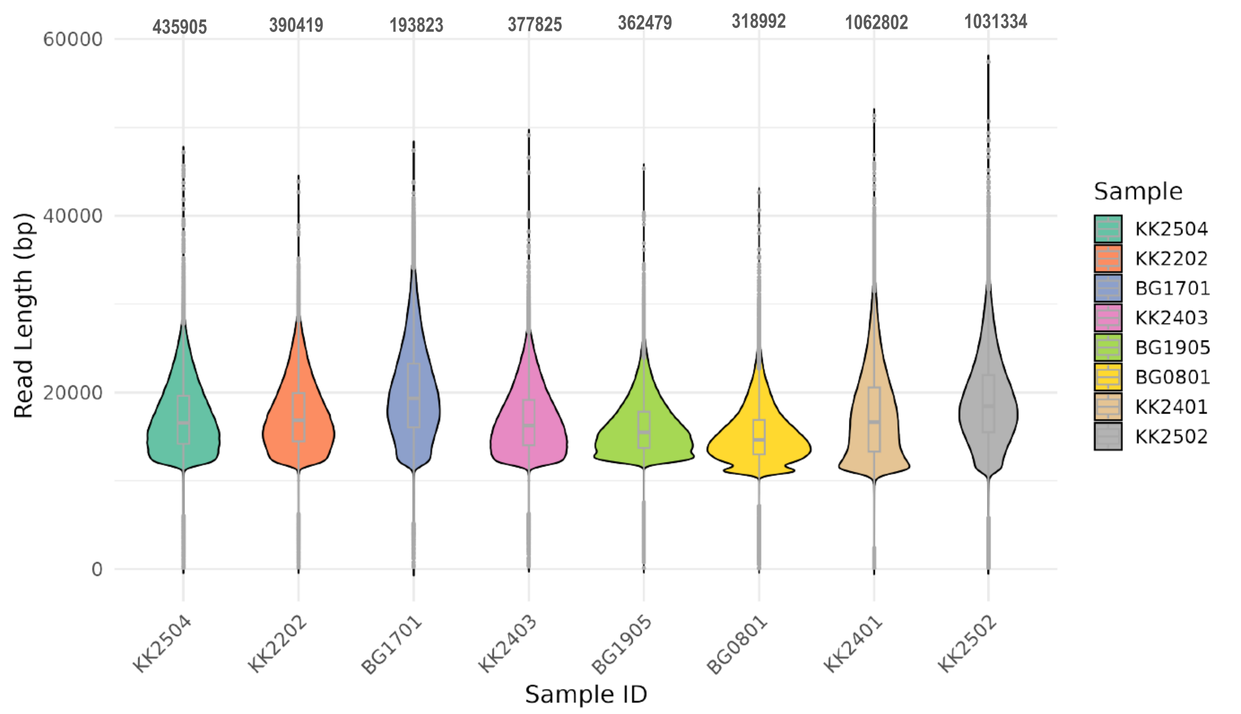

435905
390419
193823
377825
362479
318992
1062802
1031334
60000
40000
20000
0
Read Length (bp)
Sample
KK2504
KK2202
BG1701
KK2403
BG1905
BG0801
KK2401
KK2502
Sample ID


B

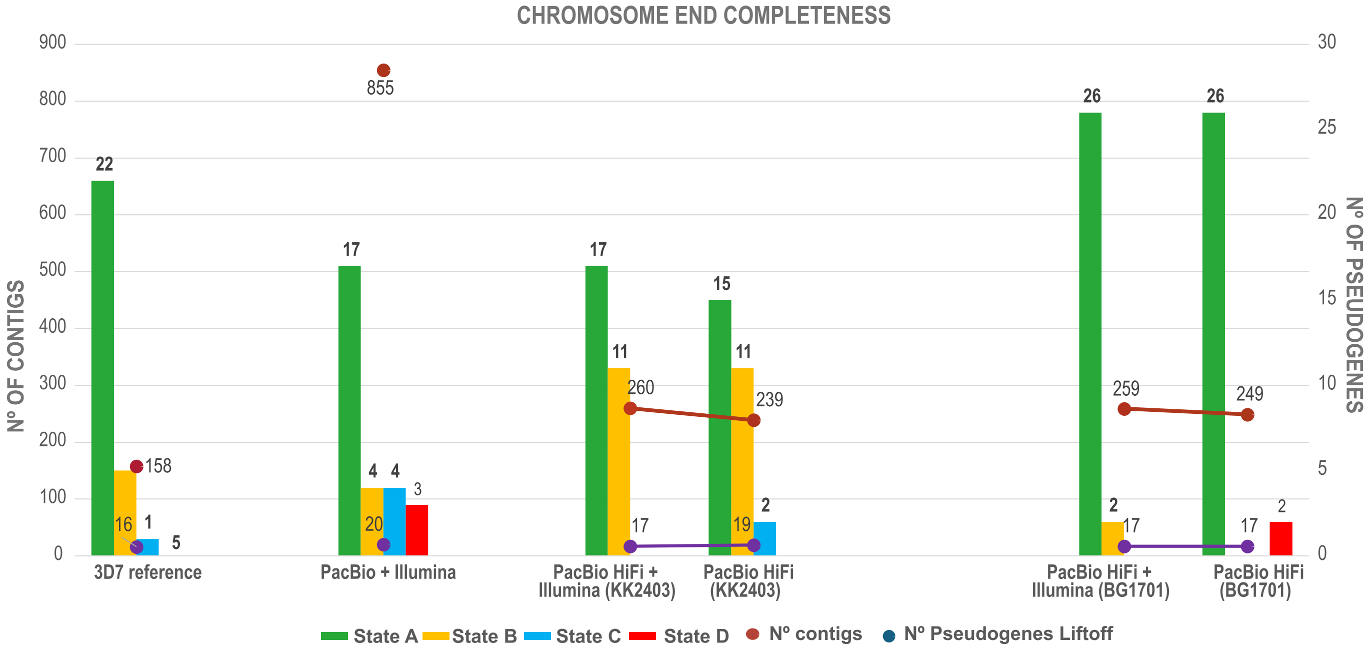

CHROMOSOME END COMPLETENESS
Nº OF CONTIGS
Nº OF PSEUDOGENES
900
800
700
600
500
400
300
200
100
0
30
25
20
15
10
5
0
22
16
1
5
158
855
17
4
4
3
20
17
11
260
17
15
11
239
19
2
26
2
259
17
26
249
17
2
3D7 reference
PacBio + Illumina
PacBio HiFi + Illumina (KK2403)
PacBio HiFi (KK2403)
PacBio HiFi + Illumina (BG1701)
PacBio HiFi (BG1701)
State A
State B
State C
State D
Nº contigs
Nº Pseudogenes Liftoff
