## Supplementary Figure 2 for "Improved long-read sequencing technology reveals full genome architecture of Malian *Plasmodium falciparum* field isolates"

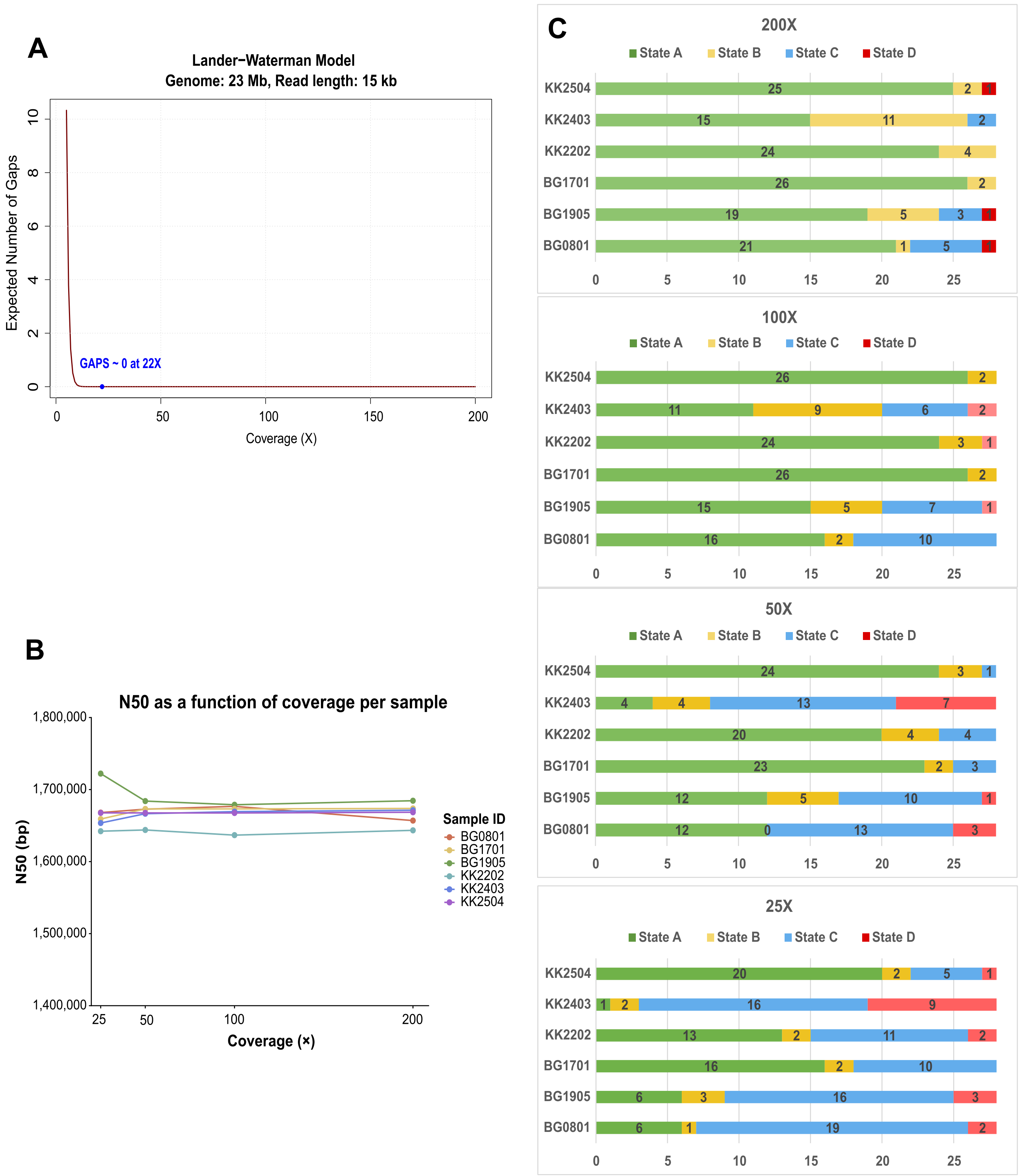
A
Lander−Waterman Model
Genome: 23 Mb, Read length: 15 kb
Expected Number of Gaps
GAPS ~ 0 at 22X
Coverage (X)
0
2
4
6
8
10
0
50
100
150
200
B
N50 as a function of coverage per sample
N50 (bp)
1,800,000
1,700,000
1,600,000
1,500,000
1,400,000
25
50
100
200
Coverage (×)
Sample ID
BG0801
BG1701
BG1905
KK2202
KK2403
KK2504
C
200X
State A
State B
State C
State D
KK2504 25 2 1
KK2403 15 11 2
KK2202 24 4
BG1701 26 2
BG1905 19 5 3 1
BG0801 21 1 5 1
0 5 10 15 20 25
100X
State A
State B
State C
State D
KK2504 26 2
KK2403 11 9 6 2
KK2202 24 3 1
BG1701 26 2
BG1905 15 5 7 1
BG0801 16 2 10
0 5 10 15 20 25
50X
State A
State B
State C
State D
KK2504 24 3 1
KK2403 4 4 13 7
KK2202 20 4 4
BG1701 23 2 3
BG1905 12 5 10 1
BG0801 12 0 13 3
0 5 10 15 20 25
25X
State A
State B
State C
State D
KK2504 20 2 5 1
KK2403 1 2 16 9
KK2202 13 2 11 2
BG1701 16 2 10
BG1905 6 3 16 3
BG0801 6 1 19 2
0 5 10 15 20 25
